# An *in vitro* platform for characterizing axonal electrophysiology of individual human iPSC-derived nociceptors

**DOI:** 10.1101/2025.01.08.631429

**Authors:** Blandine F. Clément, Lorenzo Petrella, Lea Wallimann, Jens Duru, Christina M. Tringides, János Vörös, Tobias Ruff

## Abstract

Current treatments against severe forms of neuropathic pain demonstrate insufficient efficacy or lead to unwanted side effects as they fail to specifically target the affected nociceptors - a specialized subclass of sensory neurons conveying potentially damaging stimuli information to the central nervous system. Neuropathic pain may involve different nociceptor subtypes in different patients. Tools that can distinguish nociceptive axons would enable a more targeted compound screening. Therefore, we developed an *in vitro* platform combining a CMOS-based high-density microelectrode array with a polydimethylsiloxane (PDMS) guiding microstructure that captures the electrophysiological responses of nociceptors. Human induced pluripotent stem cell-derived (iPSC) nociceptors were cultured at low density with axons distributed through parallel 4 × 10 µm microchannels exiting the seeding well before converging to a bigger axon-collecting channel. This configuration allowed the measurement of stimulation-induced responses of individual axons. Nociceptors were found to exhibit a great diversity of electrophysiological response profiles that can be classified into different functional archetypes. Moreover, we show that some responses are affected by applying the TRPV1 agonist capsaicin. Overall, results using our platform demonstrate that we were able to distinguish nociceptive axons from different subtypes. The platform provides a promising tool for screening potential candidates for nociceptor-specific drugs.

## 1. Introduction

Neuropathic pain represents a significant clinical challenge, affecting approximately 8% of the worldwide population [48]. The majority of patients do not respond adequately to existing treatments, which primarily include anticonvulsants, antidepressants or opioids. These current therapeutic approaches lack specificity and often result in adverse dose-limiting systemic side effects, such as drug dependence and tolerance [14, 45], underlining the critical need for more target-selective drugs. Neuropathic pain typically originates from dysfunctional nociceptors, a specialized subclass of sensory neurons located in the dorsal root ganglia (DRG) that transduce noxious stimuli from peripheral tissues to the central nervous system, where those signals are interpreted as pain [13]. The activity pattern and activation threshold of nociceptors are largely determined by the type and expression level of different membrane receptors and ion channels [16, 13]. Functional alterations in ion channels have been shown to influence activity profiles, [3, 27] suggesting that sensory neuron subtypes could potentially be distinguished based on these profiles [28, 31, 19, 30, 29]. Indeed, sensory neuron heterogeneity in pigs and humans has been classified through subtype-specific variations in the activity-dependent slowing (ADS) profile in which repeated stimulation increases action potential propagation time [40, 33, 35, 41]. In pathological conditions, the activity of certain nociceptors changes towards hyperexcitability which manifests in abnormal pain sensations [1, 22, 10]. The observed hyperexcitability can, among other mechanisms, again be attributed to altered expression levels or function of voltage-gated sodium channels (*e*.*g*. due to a mutation on Nav1.7), potassium channels, or specific membrane receptors [49, 2, 8]. Additionally, hyper-excitability was found to correlate with decreased ADS [34] which shows its clinical relevance [6], but also demonstrates the importance for understanding voltage-gated sodium currents [26] as promising alternative therapeutic targets to non-specific drugs [21]. Indeed, computational modelling of human C-cutaneous fiber ADS behaviors demonstrated how specific altered molecular mechanisms — including delayed sodium currents, modified ion channel conductance, and intra-axonal sodium accumulation — contribute to nociceptor dysfunction [20, 46, 37, 25]. These findings suggest the translational potential of studying ADS profiles using *in vitro* models to identify specific molecular target to a malfunctioning nociceptor sub-type and thus modulate more effectively their activity while reducing side effects. Nevertheless, ADS profile measurements of human nociceptor axons *in vitro* pose significant challenges. Two primary obstacles have emerged: 1) limited access to human nociceptors for *in vitro* models, and 2) technical challenges in recording changes in axonal membrane potentials *in vitro* due to extremely small axon diameters. Recent advancements in human induced pluripotent stem cell (iPSC) technologies have begun to address the first challenge, enabling the development of disease models for personalized pain therapeutic strategies [5, 50, 28, 31, 17]. Regarding the second challenge, despite the unprecedented precision of patch-clamp in single-cell recording, this technique is not suitable for the required experiments due to its limited repeatability and low-throughput impeding robust population-level characterization, needed for personalized medicine. Conventional higher throughput electrophysiological *in vitro* assays consist of standard DRG neuron cultures forming random and overlapping outgrowths, allow firing rate measurement as a neuronal hyperexcitability indicator [32, 17]. However, those platforms lack the precision needed to study axonal conduction properties. Recent technological innovations in microfluidic platforms have enhanced axon-specific electrophysiological investigations [23] by separating neuronal cell bodies from axons [47, 43, 42, 44] but both microscopy-based and electrophysiological read-outs lack specific axon subtype investigation. Based on existing results from the literature we hypothesized that (1) we can extract distinguishable activity response profiles by varying stimulation parameters and (2) that distributing axons into individual recording microchannels improves recording resolution by reducing interference from neighboring spike signals. Therefore we integrated polydimethylsiloxane (PDMS)-based axon guidance microstructures with high-density microelectrode arrays (HD-MEA) that enabled us to measure stimulation-induced response profiles from human iPSC-derived nociceptors (Figure 1). The specific network design allowed us to distinguish spikes from individual axons emerging from a population of diverse nociceptors. Instead of relying on spontaneous activity as a measure of excitability, the networks were exposed to various electrical stimulation paradigms for reliable ADS profile extraction and downstream characterization. By using human iPSC-derived neurons we aimed to enhance the translational relevance of our results. Overall this method provided a set of novel and comprehensive metrics about single axon responses (Figure 2B). Finally, we demonstrated the potential of the system as a drug screening platform by exposing the networks to different concentrations of the TRPV1 channel activator, capsaicin, and measuring its effects on the activity profiles of individual axons.

**Figure 1:**
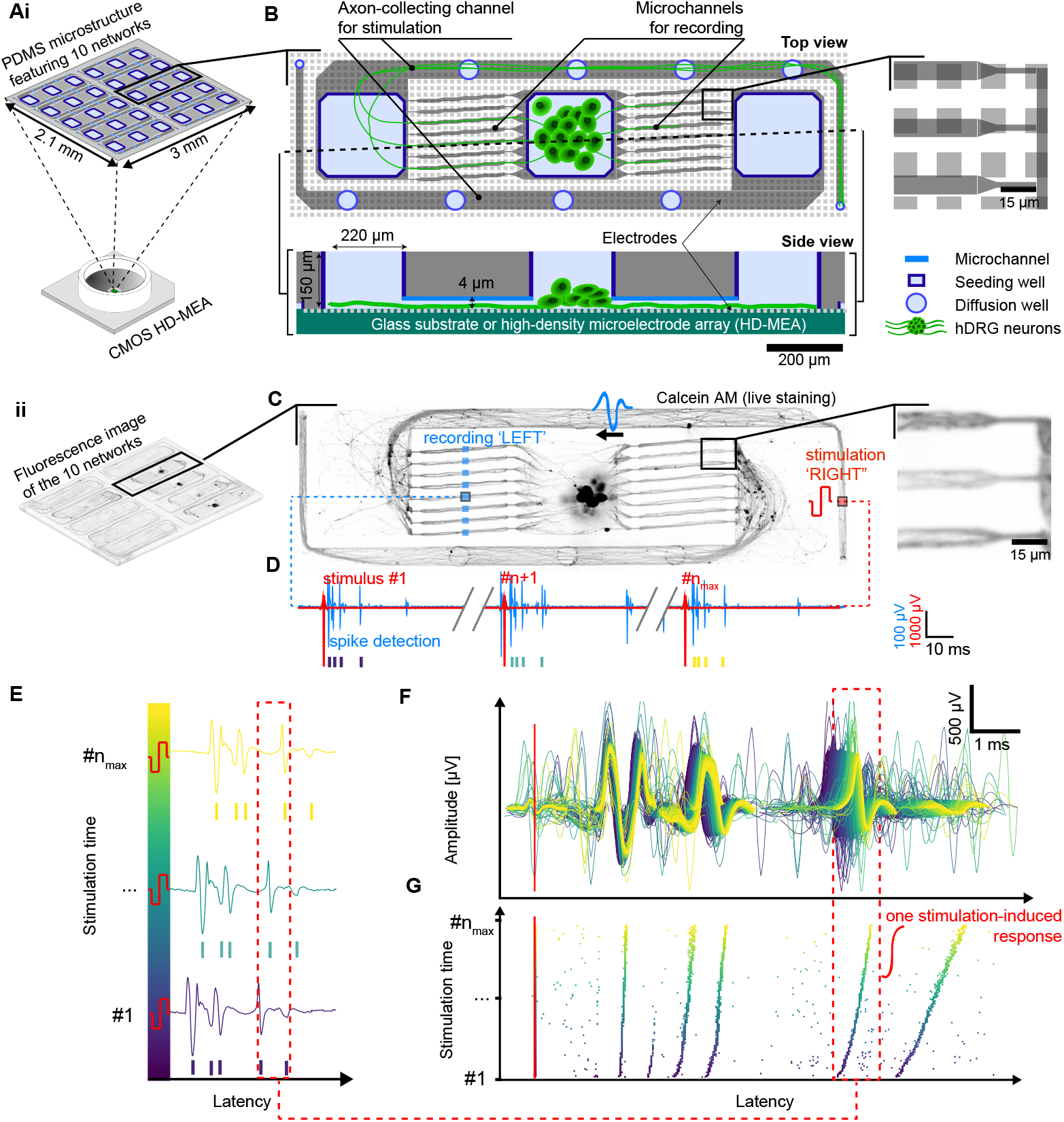
Overview of the *in vitro* platform for measuring individual nociceptor electrophysiological response on high-density microelectrode array (HD-MEA) **A**: A PDMS-based microstructure, featuring 10 identical replicates of the axon guidance network design, is placed onto a HD-MEA to measure axonal conduction properties. ii: Fluorescence image (live cells stained with calcein AM) of human iPSC-derived nociceptors (hDRG) growing inside the networks (see also Suppl. Fig. 1). **B**: Top and side view of the CAD layout of one *in vitro* axon guidance network design. Microchannels are only 4 µm tall to prevent the cells bodies from migrating inside microchannels. One seeding well in the center (open from the top) enable manual seeding of neural spheroids. Four diffusion holes along the axon-collecting channel facilitate diffusion of nutrients and oxygen and enable stimulation of axonal ends where the channel narrows down in the last segment. The high density of electrodes (light grey squares) allows for the full coverage of the network for stimulation and recording. Right subpanel shows zoom insert into the microchannels and electrode array. **C**: A hDRG spheroid was seeded in the centre seeding well of each network and grew axons that exited the seeding well through any of the 16 4×3 [h×w] µm microchannels. Within 20 days, axons subsequently merge into the 4×50 [h×w] µm axon collecting channel. Right subpanel shows a zoom into the axon microchannels. Stimulation and recording locations in the network are indicated in red and blue respectively. **D**: Example of measured voltage signal traces at the stimulation electrode (red, stimulation artefact) and at one single electrode inside a microchannel (blue) throughout the stimulation period. **E**: Three examples of the voltage signal measured by a recording electrode in a microchannel after the first (purple), x^th^ (green) and 1000^th^ (n_max_, yellow) stimulation. (The color code of the stimulation pulse number is also provided). **F**: The measured spike waveforms from one microchannel are overlayed and plotted in different color, corresponding to the time of the stimulation pulse, in order to visualize the change in spike amplitude during the 1000 stimulation cycles. **G**: The spike latencies are extracted from the voltage traces3and are overlayed along the vertical axis to visualize the change in spike latency during continued stimulation.

**Figure 2:**
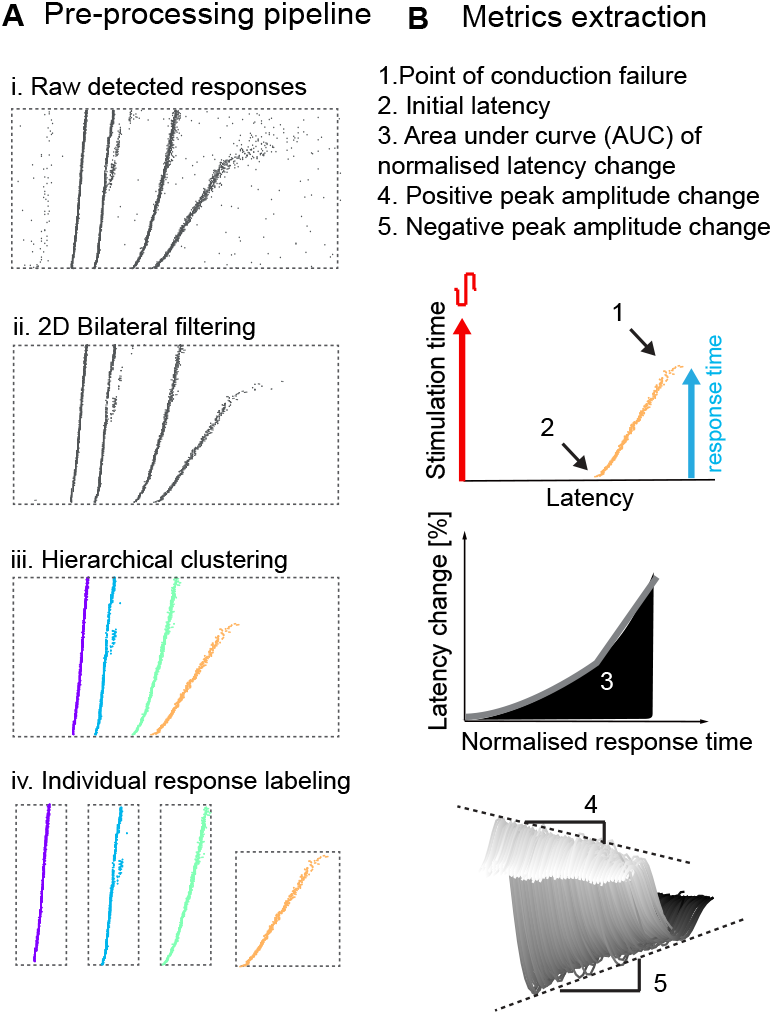
Pre-processing and analysis pipeline to characterize the different nociceptive axon responses. **A**: The different processing steps to identify the individual responses. B: Illustration of the extracted metrics to characterize the stimulation-induced responses.

## 2. Material and methods

### 2.1. Sample preparation

#### 2.1.1. MEA chip preparation

For electrophysiological recordings of *in vitro* neuron cultures, a CMOS-based HD-MEA (MaxOne, MaxWell Biosystems AG, Switzerland) was used featuring 26’400 electrodes allowing for the simultaneous recording from up to 1024 electrodes at 20 kHz. The chips were coated with 0.1 mg/mL poly-L-ornithine (Sigma-Aldrich, P4957) for at least 30 min at room temperature and subsequently washed twice with ultrapure water.

#### 2.1.2. PDMS microstructures preparation

PDMS microstructure layouts were designed using Auto-CAD (Autodesk). PDMS microstructure designs were fabricated by WunderliChips GmbH (Zurich, Switzerland). As shown in detail in Suppl. Fig 1, each network features one central seeding well separated from two lateral wells by an array of eight (per side) 10 µm-wide, 4 µm high, 300 µm-long microchannels. The channels have 3 µm-wide constrictions at both ends to minimize the number of axons that can grow through. Each lateral well is connected to one 50 µm-wide axon-collecting channel, to allow for axons to merge and grow until a designated terminal stimulation area.

PDMS microstructures were manually cut out of a wafer, exposed to isopropanol for 15 min, rinsed three times using sterile ultrapure water and dried at room temperature for at least one hour after aspirating as much liquid as possible. It is essential that both the microstructure and the MEA sensing area are perfectly dry for the placement described in the next paragraph.

#### 2.1.3. Mounting PDMS microstructure on MEA chip

Microstructures were then manually mounted onto the 3.85 mm × 2.10 mm large sensing area, making sure they were placed in the center aligned to the axes of the rectangular sensing area. The reference electrode surrounding the sensing area was left uncovered. The MEA chip was then filled with phosphate buffered saline (PBS, Sigma-Aldrich, 10010015) pre-warmed to 37 °C and desiccated to remove air pockets from within the microchannels. Microstructure placement and adhesion were then assessed by collecting an impedance map of the electrode surface as described previously in [11]. Briefly, a sinusoidal electrical signal (16 mV peak-to-peak voltage, 1 kHz frequency) was generated using an on-chip function generator. The sinusoidal signal was applied to the circumferential reference electrode of the MEA and the received signal at the microelectrodes was recorded. A highly attenuated signal indicated the electrodes covered by PDMS. Iteratively, desiccation was repeated if the impedance map showed the presence of air pockets trapped inside the microchannels. Finally, after sufficient adhesion, the PBS was replaced with a laminin coating solution consisting of iMatrix-511 (SILK, Anatomic) diluted 1:50 in DPBS(-/-, Sigma-Aldrich, D8537) for at least 1 hour or until cell seeding.

#### 2.1.4. Human iPSC-derived nociceptor spheroid formation

Human iPSC-derived nociceptors (RealDRG™ Sensory Neurons, Anatomic Incorporated) cryovial was removed from liquid nitrogen storage and was placed into a 37 °C bath. Cells were thawed (<1 minute) by gently swirling the vial in the bath until there was just a small bit of ice left in the vial. After taking the cryovial into a laminar flow cabinet, the cells were gently transferred into a sterile centrifuge tube. The cryovial was rinsed with 1 mL of DMEM/F12 (Gibco™) pre-warmed to room temperature transferring its content also to the sterile centrifuge tube. 8 mL of warmed DMEM/F12 was added slowly to the cell suspension in the tube, which was then centrifuged at 300 g for 4 minutes. After the centrifugation, the supernatant was aspirated without disturbing the pellet. The cell pellet was resuspended in 1 mL of Senso-MM Maturation Media (Anatomic Incorporated) to create a homogeneous cell suspension. After performing a cell count, the cell suspension was dispensed in one or several wells of a spheroid-forming wellplate (Sphericalplate 5D^®^, Kugelmeiers^®^), pre-filled with 1 mL of warm Senso-MM Maturation Media, with the volume necessary to create spheroids of approximately 250 - 500 cells per spheroid. The spheroid plate was centrifuged at 50 g for 3 minutes and left in the incubator overnight.

#### 2.1.5. Spheroid seeding and maintenance

Just before seeding, the diluted laminin coating solution was replaced by Senso-MM maturation medium. Spheroids were seeded between 12 to 48 hours after their preparation. One spheroid was placed in the central well of each of the 10 networks in the microstructure using a 10 µL pipette. In this process, additional care has to be put into not touching the microstructure, as it may cause its detachment from the microelectrode array. Culture media was changed a few hours after seeding. After 3 days, the maturation medium was changed to a custom-made medium and half of the medium was changed twice a week. The custom-made medium was composed of Neurobasal (NB) (Gibco ^*TM*^) medium, to which 5 % of B27 supplement (17504-044), 1 % GlutaMAX (35050-061) and 1 % Pen-Strep (15070-063, all from ThermoFisher) were added. Growth factor supplements were freshly added to 10 mL of neurobasal medium to obtain the following final concentrations: 50 ng/mL of nerve growth factor (NGF, 450-01), 20 ng/mL of brain-derived neurotrophic factor (BDNF, 450-02), 20 ng/mL of glial-derived neutrophic factor (GDNF, 450-10), 20 ng/mL NT-3 (450-03, all from PeproTech) and 5 µ M of forskolin (66575-29-9, Sigma Aldrich).

### 2.2. Experimental setup and data collection

#### 2.2.1. HD-MEA chip handling

Electrophysiological recordings were performed in an incubator with 35°C air temperature, 90% humidity and 5% CO_2_ concentration; this setup allowed for continuous recordings up to several hours. After placing the MEA chip on the recording unit, it was allowed to rest for at least five minutes before a recording session was initiated, ensuring that any spontaneous activity disrupted by movement or changes in CO_2_ levels and temperature could return to baseline.

#### 2.2.2. Network selection

Networks were selected utilizing the previously mentioned impedance map. A threshold was applied to identify electrodes not covered by the microstructure. A selection of about 1000 electrodes for each network from which to record electrical signals was performed using a custom-built previously published software [12]. Selected electrodes were then routed to available amplifiers and the resulting configuration was downloaded to the chip using the Python application programming interface (API) provided by the chip manufacturer (MaxWell Biosystems).

#### 2.2.3. Voltage recording

The network activity was tracked by recording the voltage on the routed electrodes. Voltage recordings were acquired at 20 kHz sampling frequency with a resolution of 10 bits and a recording range of approximately ± 3.2 mV, which results in a least significant bit (LSB) corresponding to 6.3 µV. Using custom software based on the MaxWell Python API, the raw traces were recorded and stored as HDF5-files.

#### 2.2.4. Voltage ramp and frequency ramp stimulation

To apply a repetitive stimulation pattern to a network, a custom Python script was created. The script makes use of the MaxWell Python API to send commands to the system hub. A biphasic pulse with a leading cathodic phase and 400 µs pulse width was used with a stimulation amplitude defined by the specific experiment. The amplitudes were rounded to the closest value available on the 10-bit DAC with approximately 3.2 mV step size. In the voltage ramp experiment, 15 networks distributed across two independent chips were stimulated and recorded at DIV 61. The stimulation, consisting in 480 pulses at 8 Hz at one specific voltage amplitude, was applied to the “LEFT” side end of axon-collecting channel of the networks one after the other, and then to the “RIGHT” side end of axon-collecting channel of the networks one after the other. Then, the next voltage amplitude was applied. The test voltage amplitudes were ranging from 300 to 1000 mV (with an incremental step of 100 mV) and the order was randomized. In the frequency ramp experiment, 20 networks distributed across two independent chips were stimulated and recorded at DIV 33 and DIV 47. Similarly to the voltage ramp protocol, the stimulation, consisting in 1000 pulses at 1000 mV pulse amplitude Hz at one specific frequency, was applied to the “LEFT” side end of the networks one after the other, and then applied to the “RIGHT” side of the networks one after the other. The test stimulation frequencies were the following: 1 Hz, 2 Hz, 4 Hz, 8 Hz, 16 Hz, 32 Hz and 64 Hz, and their order was randomized. Each single network rested for at least 10 minutes between consecutive stimulation trains in both protocols.

#### 2.2.5. Capsaicin stimulation

The tested capsaicin concentrations were selected based on values reported in [36]: 100 nM, 1 µM, and 10 µM. To achieve these concentrations, 1 g of lyophilized capsaicin (Sigma Aldrich, PHR1450-1G) was initially dissolved in 3.273 mL of dimethyl sulfoxide (DMSO) resulting in a 1 M capsaicin solution. All capsaicin solutions were prepared such that the DMSO concentration did not exceed 0.001%. Then, 1 µL of the DMSO-capsaicin solution was added to 10 mL of 37 °C NB media resulting in 0.1 mM solution. It is worth noting that the DMSO-capsaicin solution was added by slowly releasing the liquid into the media while moving the pipette around in the solution. The warming and slow release helped to ensure that the DMSO-capsaicin solution fully dissolved and did not stick to the sides of the falcon tube, due to the low solubility of the capsaicin. This solution was then further diluted by adding 1 µL, 10 µL or 100 µL in respectively 999 µL, 990 µL and 900 µL of warm NB media to achieve the respective testing concentrations. All three solutions were first warmed up in the bath before use in the chip chamber. The experiment consisted in replacing the chip chamber medium volume by the test capsaicin solution, to control any effect due to a full medium change or medium evaporation over measurement time. Additional control consisted in performing a full medium change. After replacing the solution with the next one, the sample was put back in the incubator, and the stimulation protocol (1000 pulses, voltage amplitude of 1000 mV, 32 Hz) was applied 5 minutes after the CO_2_ level in the environment of the incubator went back up to 5%. For washout, the chip was rinsed 5 times, every rinse consisted in removing 900 µL and replacing by 900 µL.

#### 2.2.6. Image acquisition and analysis

An inverted confocal laser scanning microscope (FluoView 3000, Olympus) was used to image the fluorescently labeled cultures. Two channels were typically acquired: 488 nm (Calcein AM or CMFDA) and phase contrast brightfield images (only for glass bottom dishes). Fluorescence imaging of the CMOS chips was done by removing all excess medium and using a round glass coverslip (10 mm diameter) mounted on top of the PDMS microstructure. The surface tension between the cover slip and the chip enabled us to invert the chip for imaging in an inverted microscope. For mounting in the CLSM, the CMOS chip was placed in the recording unit which in turn was mounted into a custom made metal insert that fits into the stage of the microscope. This configuration enabled inverted mounting for imaging [11]. Microscope images were processed using Fiji [39]. Importantly, due to their size, stained soma are brighter and thus more visible than axons on microscopy images. To enhance the intensity of the axons compared to the soma, a pixel logarithm operator was applied to all the representative fluorescent images shown in the figures of this paper (raw image also show in Suppl. Fig. 2). The brightness and contrast were manually adjusted to suppress background fluorescence.

### 2.3. Data processing and analysis

### 2.4. Raw data processing

Raw data were processed using a band-pass filter (4th order acausal Butterworth filter, 300-3500 Hz). The baseline noise of the signal was characterized using the median absolute deviation for each electrode [38]. Action potential event times were extracted from the voltage trace by identifying negative signal peaks below a threshold of 10 times the baseline noise. Successive events within 0.75 ms were discarded to avoid multiple detection of the same spike. Spike amplitudes were defined as the absolute value of the negative amplitude of the detected peak. Spike waveforms were extracted from the filtered voltage trace using the data within a -1 ms to 1 ms window around the timestamp of the detected spike. The waveforms were also used to measure the spike amplitude.

#### 2.4.1. Individual microchannel identification

Every single microchannel is physically spanned by a set of electrodes, which results in row-like arrangement of pixels on the impedance map representation of the microstructure. Therefore, microchannels were visually identified on the impedance scan and respective electrodes/pixels were manually selected via a custom-made GUI [12] and saved in a separate file.

#### 2.4.2. Stimulation-induced response analysis

Spike-triggered waveform and raster plots were generated for each electrode (one per microchannel) to visually inspect the response of axons in each microchannel to the various stimulation protocols. As the stimulation signal saturates the amplifiers (clamped at 3.2 mV) independently from the stimulation amplitude, an artefact amplitude threshold of 3 mV was established to detect the stimulation pulse times on the stimulation electrode. Then, for the most central active electrode of each microchannel, spikes were detected within 20 ms from the stimulation artefact (but only 15 ms in case of the 64 Hz stimulation due to the resulting inter-stimulation interval). The resulting spike waveform and latency raster plots are presented in Fig.1G and H, respectively. This time window was sufficiently long for the action potentials to travel from the stimulation electrode to the microchannels.

#### 2.4.3. Identification of individual responses

The raster plot that contains the time of each recorded spike is a 2D array image in which the spontaneous random spikes correspond to background noise and the stimulation induced spikes form “bands” which are the features of interest. In order to remove the background noise, a bilateral filter was applied (Python OpenCV library). A clustering algorithm (hierarchical clustering, Python scikit-learn library) was then applied to separate and identify the “bands”. It is important to note that the metrics extraction is affected by the band detection method, and that specific corner cases arise and have a impact on the variability. In the voltage ramp analysis, responses originating from the same axon across multiple stimulation voltages were identified by their matching latency values with a tolerance of 0.2 ms to account for small latency shifts. At each stimulation voltage newly appearing bands were identified as those not present at the previous lower stimulation voltage.

#### 2.4.4. Metrics extraction

Response characteristics were quantified using multiple metrics as shown in Figure 2. We assessed conduction failure by identifying the final stimulation pulse that elicited a detectable response. The ratio of the time until the final elicited response over the total duration of stimulation defined the “point of conduction failure”. The conduction speed was calculated from the initial latency—defined as the time delay between the first stimulus and detecting the induced response at the recording site. Following the approach of Dickie et al. [9], we characterized the progressive slowdown in conduction by calculating the area under the curve (AUC) of normalized latency changes throughout the normalized response period. We decided to normalise both dimensions so that the AUC value would not be impacted by the late latency or the length of the response, but would only reflect the bending behavior of the “band”. For each detected spike event, the percentage change in latency was calculated according to the following equation:

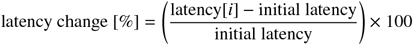

where latency[*i*] represents the latency at stimulation iteration *i*. The response period was normalized by dividing the stimulus repetition numbers by the last stimulus number for which a spike event was detected, resulting in an array of values from 0 to 1. Changes in waveform characteristics throughout the stimulation period were assessed by performing linear regression analysis on both negative and positive peak amplitudes across successive stimulation pulses. The resulting slopes define the changes in peak amplitude. To account for response variability, we included the coefficient of determination (R-value) from these fits as a measure of the linearity. While extracellularly recorded action potentials in slowly conducting nerve fibers typically have a triphasic shape (2 positive peaks, 1 large negative peak) in pigs [46], the positive peak amplitude we report always corresponds to the first positive peak only.

#### 2.4.5. Cluster analysis of the different electrophysiological profiles

Electrophysiological responses were recorded from 320 microchannels distributed across 20 networks on two chips. Each axon-collecting channel per network side was stimulated with 1000 pulses at 16 Hz with an amplitude of 1000 mV. Multiple metrics characterizing the elicited responses were extracted from these recordings as described in 2.4.4. Prior to analysis, some responses were filtered using specific exclusion criteria. We eliminated responses that initiated past 20% of the total stimulation period, as these likely resulted from event misdetection due to temporal proximity of adjacent events (see Suppl. Fig. 6 for all corner cases and sources of error in response identification). For dimensional reduction, principal component analysis (PCA) was performed on the scaled feature space. The number of retained principal components was determined by setting a threshold of 0.75 for cumulative explained variance. Cluster analysis was then performed using the K-means algorithm. To determine the optimal number of clusters, silhouette scores were calculated for cluster numbers ranging from 2 to 10, and the configuration yielding the highest silhouette score was selected for the final clustering analysis in the PCA space. To characterize representative responses for each cluster, we identified cluster centroids and selected the three data points closest to each centroid based on Euclidean distance in the principal component space. For these representative points, we plotted both the latency responses and corresponding waveforms to visualize typical response patterns within each cluster.

## 3. Results and discussion

### 3.1. Axon guidance structure design for measuring stimulation induced nociceptor activity from individual axons on HD-MEAs

We developed an *in vitro* platform with the intention to measure the electrophysiological response of individual nociceptor axons on a high-density microelectrode array (HD-MEAs). A polydimethylsiloxane (PDMS)-based axon guidance microstructure was designed to fit 10 identical axon guidance networks onto a single HD-MEA (see Fig. 1A). Each axon guidance structure consisted of three seeding wells: one central seeding well and two lateral seeding wells. The central seeding well contains the hDRG neurons consisting of human induced pluripotent stem cell (iPSC)-derived nociceptors (Fig. 1B). From the central seeding well, eight channels extend outward on each side and narrow down to about 3 µm before exiting into the neighboring seeding wells with the goal to reduce the probability of axonal backgrowth. Afterwards the axons converge into a wider channel that collect all axons, which narrows down in the last segment in order to confine more axons and thus improve the stimulation efficiency. Holes along the axon-collecting channels were added to improve the diffusion of nutrients, oxygen and added compounds for drug screening applications (Fig. 1B). Each of the 10 networks is point-symmetric. Within 20 days of seeding neural spheroids extend axons through the 16 axon microchannels and extend towards the end of the axon-collecting channel (Fig. 1C). To induce activity and measure axonal spike latencies, the networks were repeatedly and independently electrically stimulated at the end of each of the two axon-collecting channels using different stimulation voltages and frequencies and the corresponding spike signals were recorded within the 8 axon channels Fig. 1C, G. Thus each network was stimulated once from each side and spikes were only recorded from the 8 corresponding axon channels. Extracted spike latencies were stacked along the vertical axis to visualize any change over time (Fig. 1H). Overall, this platform supports electrophysiological analysis of up to ten individual nociceptor networks on a single high-density microelectrode array, which increases experimental throughput and strengthens the statistical reliability of the latency measurements.

### 3.2. A voltage-dependent response is obtained when stimulating the end of axon-collecting channel

As discussed previously, while the axonal initial segment is known to be the most excitable site for neuronal stimulation [7], stimulating distal axon ends better emulates biological sensory inputs from the periphery. We applied an increasing stimulation voltage at the terminal ends of the axons contained in axon-collecting channel via one stimulating electrode as illustrated in Fig. 1C and according to the protocol described in the methods. This led to evoked action potentials that propagated towards the cell somata in the central seeding well. The device geometry enabled the characterization of the response spikes independently in eight recording microchannels. As illustrated in the stimulation-latency plot example for one microchannel (Fig. 3A), the application of voltage stimulation pulses with increasing amplitudes ranging from 300 mV to 1000 mV revealed distinct response patterns. Figure 3A demonstrates that no responses were observed at 300 mV, while initial but un- sustained responses emerged at 500 mV. In contrast, stimulation at 700 mV consistently elicited six distinguishable responses, suggesting the presence of six axons in this microchannel, reproducibly recruited at 900 mV. The totality of elicited responses for two stimulated networks are shown in Suppl. Fig.

**Figure 3:**
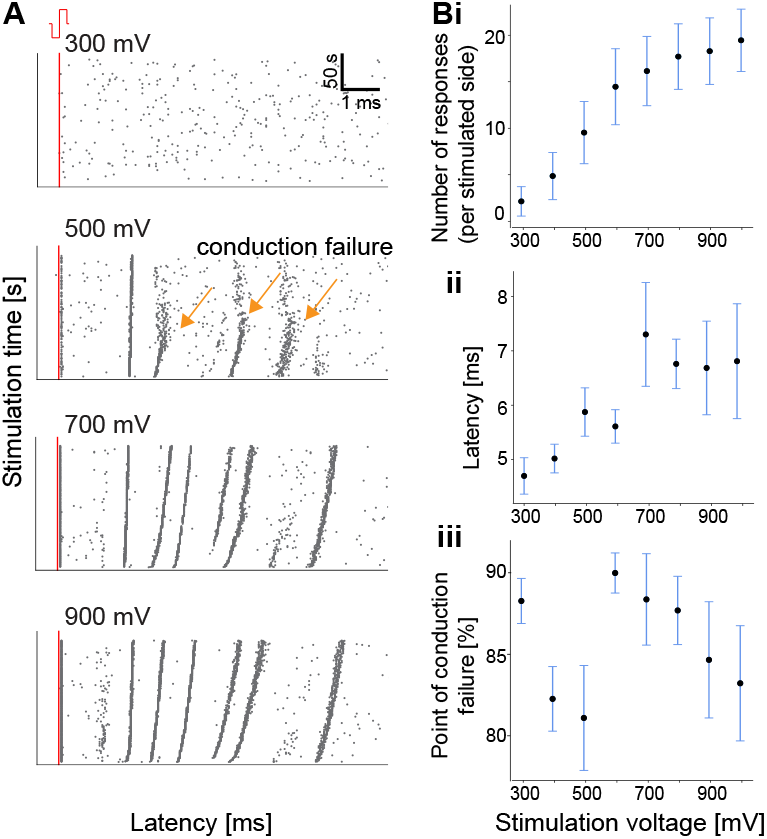
Stability of stimulation-induced responses upon varying stimulation voltage values. **A**: Representative example of elicited responses in one microchannel upon increasing stimulation voltage. **B**: Quantification of different network response parameters depending on the stimulation voltage for a total of 200 microchannels from 25 network stimulated sides spread on two chips and measured at DIV 61. Means across networks and standard error of means are plotted. i: The number of newly appearing responses increases with an increased stimulation voltage. ii: Mean latency for new elicited responses per voltage value. iii: Point of conduction failure as the percentage of the total stimulation time, for new elicited responses per voltage value. All data in B is presented as mean ± SE of mean

3. The quantified data in Fig. 3Bi shows that the number of stimulation-induced responses increases with increasing stimulation amplitudes until reaching a plateau at 800 mV, suggesting progressively increasing number of axons being recruited within the axon-collecting channel. The maximum number of about 25 stimulated axons on average indicates that a substantial portion of neurons (over 10%) grow axons that reach the stimulation site. This number is probably limited by the 3× 4 µ m cross-section of the constriction that restricts the passing of more axons. The observed voltage dependency is consistent with previous studies indicating that higher voltages are required to recruit axons located further from the electrode and those with smaller diameters [15]. Supporting this interpretation, Fig. 3Bii reveals that the mean latency of newly appearing responses gradually increases with voltage, aligning with the known relationship between axon diameter and conduction velocity [18], if we consider mean latency as a surrogate metric for conduction speed in this platform. Indeed, this platform assures that axons take the same path and grow approximately the same length from the recording site to stimulation site which enables us to measure response latency to a stimulus with sufficient precision and sensitivity to distinguish axons with different propagation speed. An interesting pattern emerges in the analysis of conduction failure. As evident from Figure 3Biii, stimulation with voltages below 600 mV results in earlier conduction failure compared to higher voltages. This observation is consistent with previous findings regarding the decreased excitability of repeatedly activated axons [6], particularly in nociceptors, where sustained activation can lead to reduced excitability patterns such as activity-dependent slowing (ADS) [46]. While the excitation threshold becomes too high for 500 mV stimulation, 700 mV is sufficient to maintain response throughout the stimulation period. However, interestingly, the point of conduction failure tends to decrease again with increasing stimulation voltage amplitudes. We could hypothesize that this effect is due to the fact that higher voltage amplitudes recruit thinner axons corresponding to a subtype that tends to exhibit earlier conduction failure. These findings suggest that achieving reliable axon recruitment at distal axon endings requires careful consideration of stimulation voltage thresholds, as lower voltages may be insufficient to maintain consistent axon activation during repeated stimulation.

### 3.3. Stimulation frequency dependence of axonal conduction

As discussed previously, nociceptors exhibit characteristic activity-dependent slowing (ADS) profiles, extensively documented in both human and porcine *in vivo* studies [35, 4]. In order to investigate whether similar behavior could be observed in this *in vitro* platform, we examined the electrophysiological responses of cultured networks to varying stimulation frequencies; and characterized the ADS profiles of individual axons to compare them with known *in vivo* patterns. In general, the results demonstrate that cultured axons exhibit frequency-dependent modulation of their conduction properties, though with some distinct differences from *in vivo* observations. As shown in the stimulation-latency plot example for one microchannel (Figure 4A) or for all microchannels of one full network (Suppl. Fig. 4), axonal responses exhibit a “bending” behavior throughout stimulation, which is quantitatively supported by the systematically increasing area under the latency change vs. normalized time curve (AUC) with increasing stimulation frequency 4Bi. This means that during repetitive stimulation, the axons tend to conduct subsequent action potentials progressively slower, with the decrease in conduction latency being more at higher stimulation frequencies. This frequency-dependent modulation becomes visible from stimulation frequencies as low as 4 Hz. This increase in latency throughout repeated stimuli is often accompanied by conduction failure, characterizing the point from which the axon stops responding to the stimulus 4Bii. Interestingly, the observed changes in response latencies were often accompanied by a progressive decrease (or more seldom increase) in negative and positive peak amplitudes of detected associated waveforms throughout such repetitive stimuli (Fig. 4Biii). Changes in negative and positive peak amplitude were also variable and the consistency of their change is represented by the coefficient of determination R (see Fig. S5 in SI). While changes in latency and conduction failure are consistent with previous *in vivo* microneurography [35] and single-neuron patch clamp studies [27], there are notable differences in the observed frequency ranges that elicit ADS. Specifically, while *in vivo* studies typically report ADS effects at lower frequencies (0.5-4 Hz), our platform showed pronounced effects at higher frequencies. This disparity may be attributed to differences between *in vitro* and *in vivo* environments, the size and composition of nociceptors in culture, and hypothetically the absence of the periaxonal space usually provided by non-myelinating Schwann cells *in vivo* which was reported as a major contributor to ADS in modelling studies [46]. The consistent measurement of peak amplitudes comes as a novel observation *in vitro* that has only been reported in computational model predictions in the form of decreased sodium channel current [46]. Despite these frequency-range differences, the fundamental ADS behavior was consistently observed across multiple preparations. However, this frequency-dependent modulation was not uniform across all recorded axons, indicating heterogeneity in their conduction properties, and confirming the potential of this platform to distinguish among different behaviors. The variation in frequency sensitivity, while noteworthy, does not diminish the platform’s ability to capture key physiological properties of axonal conduction.

**Figure 4:**
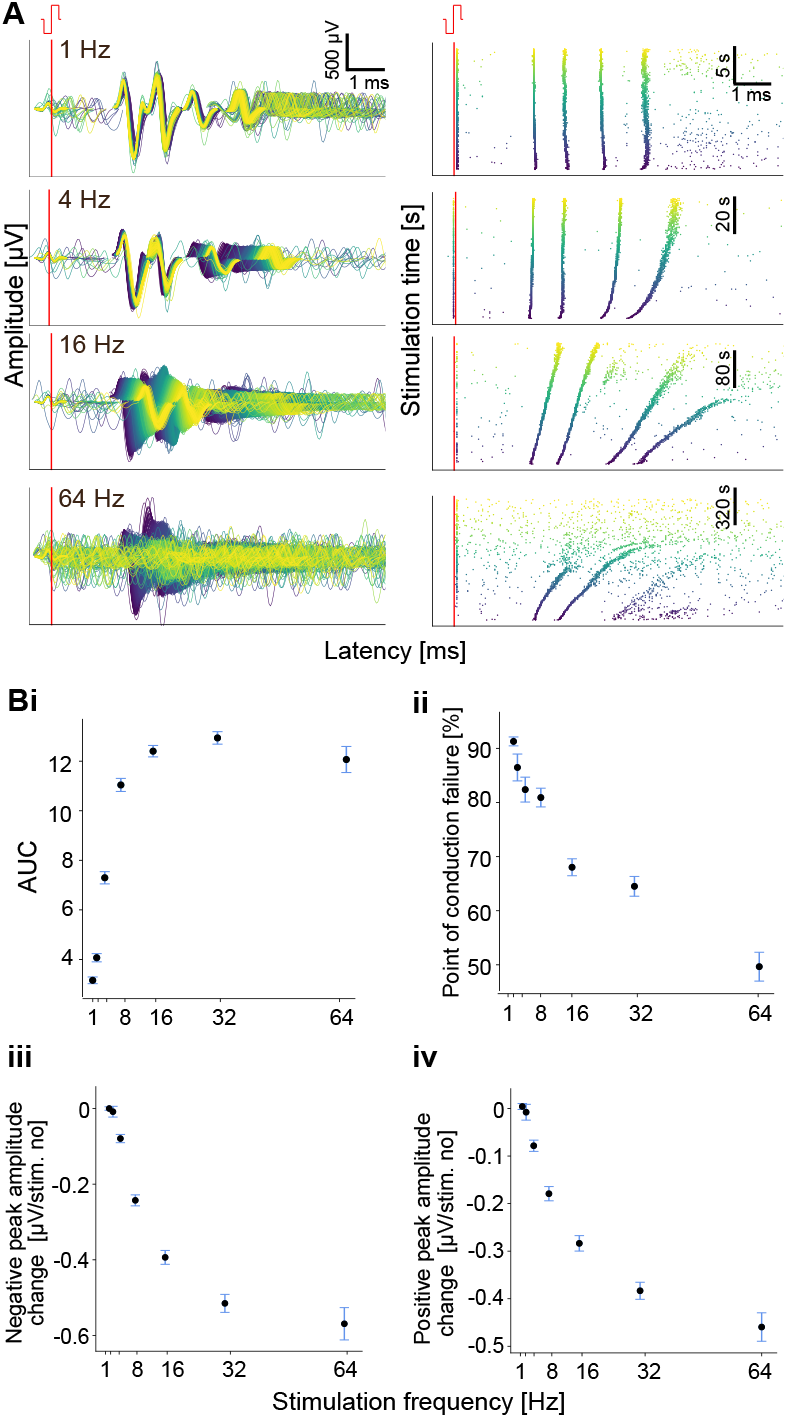
Frequency-dependent responses. **A**: Representative example of elicited responses in one microchannel upon increasing stimulation frequency. **B**: Quantification of different network parameters depending on the stimulation frequency from 20 recorded networks distributed across two chips. Each data-point represents the average from all responses from 8 microchannels of a single network side. Means across all networks sides and standard error of means are plotted. i: The change in spike latencies during the stimulation period was quantified as area under curve (AUC) and increases with increasing stimulation frequency. ii: The response conduction failure decreases with increasing stimulation frequency. iii: The change in the waveform negative peak amplitude increases with increasing stimulation frequency. iv: The change in the waveform positive peak amplitude increases with increasing stimulation frequency. All data in B is presented as mean ± SE of means

### 3.4. Axons can be clustered based on their electrophysiological response to stimulation

The present study aimed to identify potential electrophysiological signatures that could distinguish between nociceptor subtypes. As described in the introduction, the classification of nociceptors has traditionally been based on molecular markers, but functional characteristics might provide additional insight into their diversity, at population level. Generally speaking, our results reveal considerable heterogeneity in axonal conduction properties across the recorded nociceptors, suggesting potential functional distinctions between subtypes. We summarized electrophysiological responses to a given stimulation in a set of metrics capturing changes upon repetitive stimulation in axonal conduction, excitability and spike waveform, but also considering the initial latency as a potentially inherent property of nociceptors. For this analysis, we have picked the results obtained at stimulation frequency of 16 Hz because these showed many stable bands with interesting features that could be extracted using our pre-processing and analysis pipeline (Fig. 2). As shown in Figure 5Ai-v, the responses demonstrated marked variability. The AUC, which reflects the degree of conduction slowdown, and the point of conduction failure have particularly wide distributions. We employed principal component analysis (PCA) followed by cluster analysis to investigate if different axon subtypes could be identified. Using a threshold of 0.8 for cumulative explained variance resulted in three principal components (see Figures 5B and Suppl. Fig. 7 Using silhouette scoring for determining the number of clusters identified three and four clusters as optimal for our dataset, respectively scoring 0.41 and 0.39. While three clusters scores higher, we considered four clusters to maximize distinct electrophysiological profiles. These clusters are shown in distinct colors and their respective centroids are designated by the red cross. As illustrated in Figure 5C, we visualized these typical cluster profiles by plotting the three closest responses to each cluster centroid, effectively providing archetypal response patterns for each identified group. Indeed, the first identified cluster in blue seems to exhibit consistently sustained response throughout the whole stimulation, with low ADS and variable change in peak amplitudes. On the contrary, profiles highlighted in green and purple clusters exhibit earlier conduction failure. The green and purple clusters are better discriminated along the PC2 which features high weights for the two amplitude change associated metrics, suggesting that they mainly differ from each other by their change in peak amplitude throughout the stimulation period. The last identified cluster exhibits more pronounced ADS associated with conduction failure, while conserving peak amplitude throughout the response, associated with a late initial latency, suggesting a slower baseline conduction speed. This finding confirms the trend we observed in Fig. 3Biii where higher voltage amplitudes recruited axons displaying earlier conduction failure. As higher voltage amplitudes recruit thinner axons, that also conduct signal slower, thus it is consistent to observe that late latency responses also exhibit earlier conduction failure, as represented by this last archetype 4. While those representative examples provide qualitative interpretation, they demonstrate distinct conduction properties that could potentially correspond to different functional subtypes of nociceptors. It is important to acknowledge several limitations in our analysis. We used silhouette scoring to look for four clusters, but alternative clustering metrics might yield different optimal cluster numbers. Future work should validate these findings with larger datasets and perhaps incorporate additional electrophysiological parameters.

**Figure 5:**
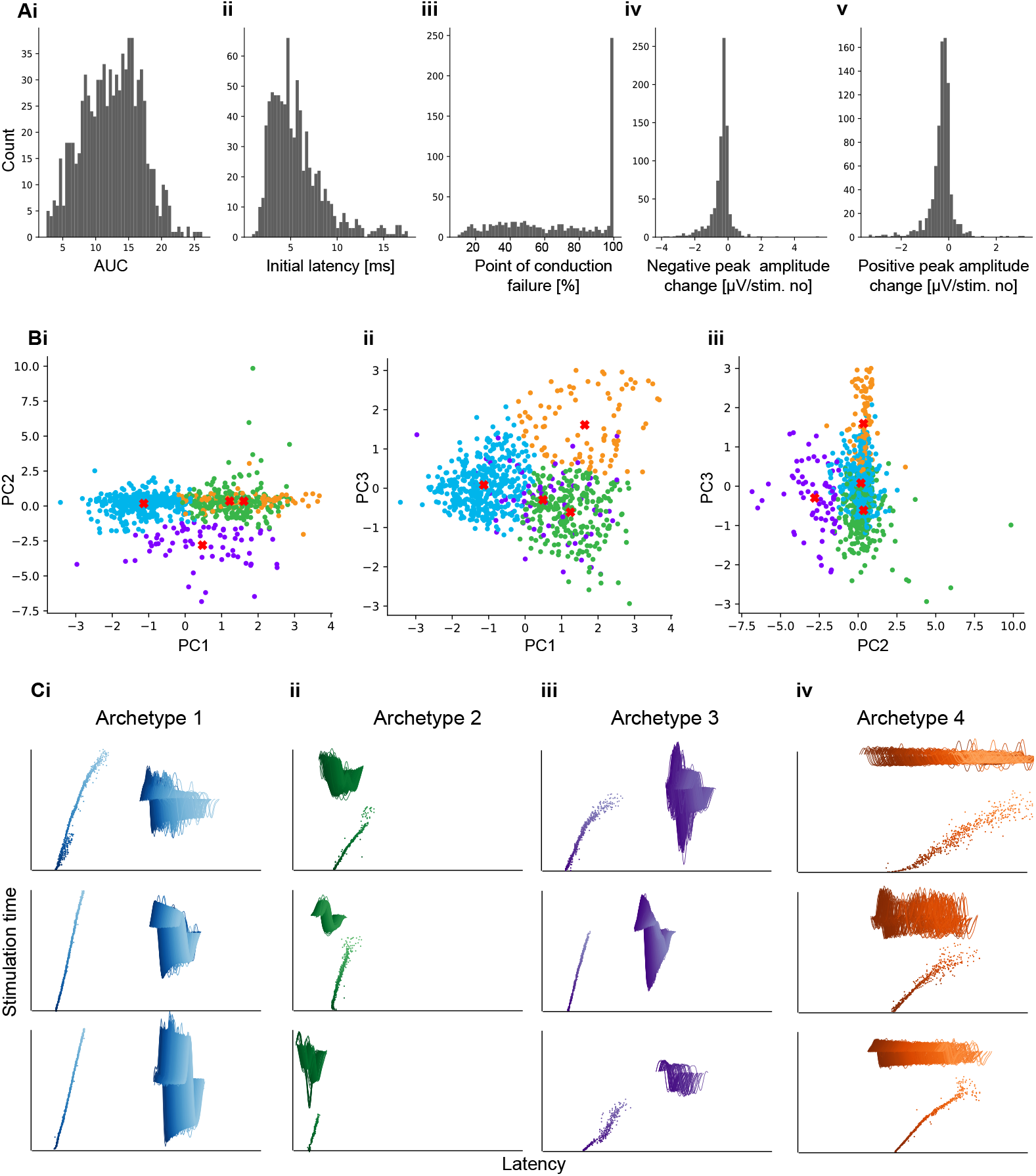
Clustering analysis of electrophysiological responses reveals different archetype responses. **A**: Distributions of each metric that was used to generate the original feature space used in the subsequent cluster analysis. Distributions summarize all individual responses to electrical stimulation (16 Hz) for a total of 320 microchannels across 20 networks distributed on 2 chips and respectively recorded at DIV 33 and DIV 47. **B**i-iii: Response data points are projected on pairs of principal components (PC) resulting from the principal component analysis (PCA) performed on the original feature space. Number of PCs was chosen using a threshold of 0.8 for the cumulative explained variance. Four clusters were identified based on silhouette scoring method (silhouette score = 0.39). **C**: Four archetype responses are illustrated by the visual representation of the stimulation-induced response combined with the corresponding action potential waveforms) corresponding to the centroid of each identified cluster.

### 3.5. Capsaicin induces a change in response in specific individual axons

As discussed in the introduction, TRPV1-positive nociceptors are specifically activated by capsaicin, providing an opportunity to investigate functional differentiation between nociceptor subtypes. Since *in vitro* neural response is highly dependent on the environmental parameters [24], we employed a controlled media change protocol to isolate capsaicin-specific effects from those that might arise from changes in ionic concentrations (See methods 2.2.5 for the details.) Our results revealed distinct response behavior among axons within the same microchannel following capsaicin administration, suggesting the presence of both capsaicin-sensitive and capsaicin-insensitive types. As shown in Figure 6, we measured three distinct axon responses to electrical stimulation at baseline in one microchannel: two responses are sustained throughout the stimulation period (green arrows), while the last response exhibit early conduction failure (orange arrow). Upon addition of capsaicin, the latest-latency response (orange arrow) shows different behavior from its baseline response while leaving the other two responses unaffected compared to their baseline. Specifically, the corresponding axon that initially displayed a conduction failure showed more consistent activation following capsaicin exposure, suggesting a sensitization effect. Indeed, capsaicin lowers the excitability threshold of TRPV1-positive nociceptors. However, at capsaicin concentrations exceeding 100 nM, this same axon reverted to a more variable response pattern. The transition from sensitization to variable response at higher concentrations could be interpreted as a shift from sensitization to desensitization, as reported by other researchers in the field. It is noteworthy that the two faster-conducting responses remained unaltered throughout all experimental conditions, suggesting these axons were TRPV1-negative. One limitation of our study was the incomplete recovery of the capsaicin-sensitive response following washout, which may indicate long-term desensitization. Nevertheless, this limitation does not diminish the primary finding that our platform can reliably distinguish between TRPV1-positive and TRPV1-negative nociceptors. These results demonstrate the potential of our platform to identify and characterize specific nociceptor subtypes based on their functional responses. Importantly, this approach offers advantages over traditional *in vitro* studies by providing more nuanced readouts beyond simple firing rate changes, potentially enabling more detailed classification of nociceptor subtypes based on their functional characteristics.

**Figure 6:**
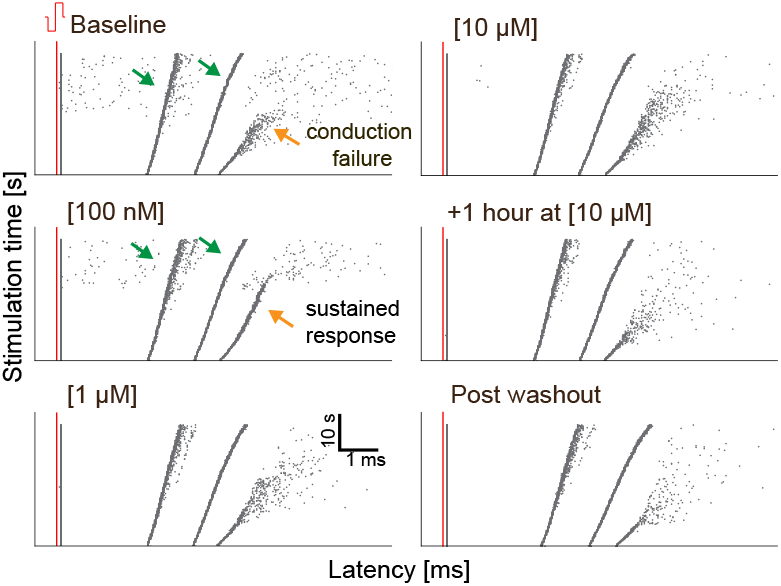
Differential responses are observed to the combined capsaicin and electrical stimulation recorded in one microchannel. Three stimulation-induced responses are detected at baseline, among which two of them seem unaffected by capsaicin addition at different concentrations (green arrows), while one response exhibits a different behavior (orange arrow).

## 4. Conclusions

Here we described an *in vitro* platform on high-density CMOS MEAs that enables the precise recording of electrophysiological response profiles from human iPSC derived nociceptors. The clear separation of stimulation and recording sites using PDMS axon guidance structures allowed us to measure stimulation-induced spike latencies. The resulting latency profiles and changes in that emerged during repetitive stimuli were used to distinguish nociceptor subtypes. In future studies validation of the functional archetypes with genetic expression profiles might establish the functional response profiles as independent biomarkers to distinguish nociceptor subtypes. Moreover, we have shown that the application of the TRPV1 channel activator capsaicin affects the extracted spike metrics. Thereby we demonstrate the potential of the presented application for future drug screening applications. When combined with automated media exchange systems [24] it might become possible to create highly reproducible, cell subtype specific dose response curves. Finally, the wide range of electrophysiological responses observed in this study highlights the need for new *in vitro* methods that offer high-resolution readout of distinct nociceptor subtypes at the individual axon level, enabling subtype specific drug discovery processes. This platform could also have applications in other areas of neurological drug research where success rate and specificity of currently used drugs are low.

## 5. Acknowledgments

This research was supported by ETH Zürich, the Swiss National Science Foundation (SNSF) (project number 182779) and the Human Frontier Science Program (HFSP) postdoctoral fellowship.

## 6. Author contributions

Conceptualization: B.F.C.,J.V, T.R.; Data Curation: B.F.C.; Formal Analysis: B.F.C., L.P., L.W.; Funding Acquisition: T.R., J.V.; Investigation: B.F.C., L.P., L.W.; Methodology: B.F.C., J.D., T.R., C.M.T; Project Administration: B.F.C., J.V., T.R.; Resources: J.V.; Software: B.F.C., J.D.; Supervision: B.F.C., J.V., T.R.; Validation: B.F.C.; Visualization: B.F.C., T.R.; Writing – Original Draft: B.F.C., T.R. C.M.T., J.D., J.V., L.P., L.W.; Writing – Review and Editing: B.F.C., T.R., C.M.T., J.D., J.V., L.P., L.W. All authors have read and agreed to the published version of the manuscript.

## Appendix A.

### Supplementary figures

Supplementary data associated with this article can be found in the online version here

## Notes

### Competing Interest Statement

The authors have declared no competing interest.

