## Supplementary Information for "An *in vitro* platform for characterizing axonal electrophysiology of individual human iPSC-derived nociceptors"

### S1 Supplementary Figures

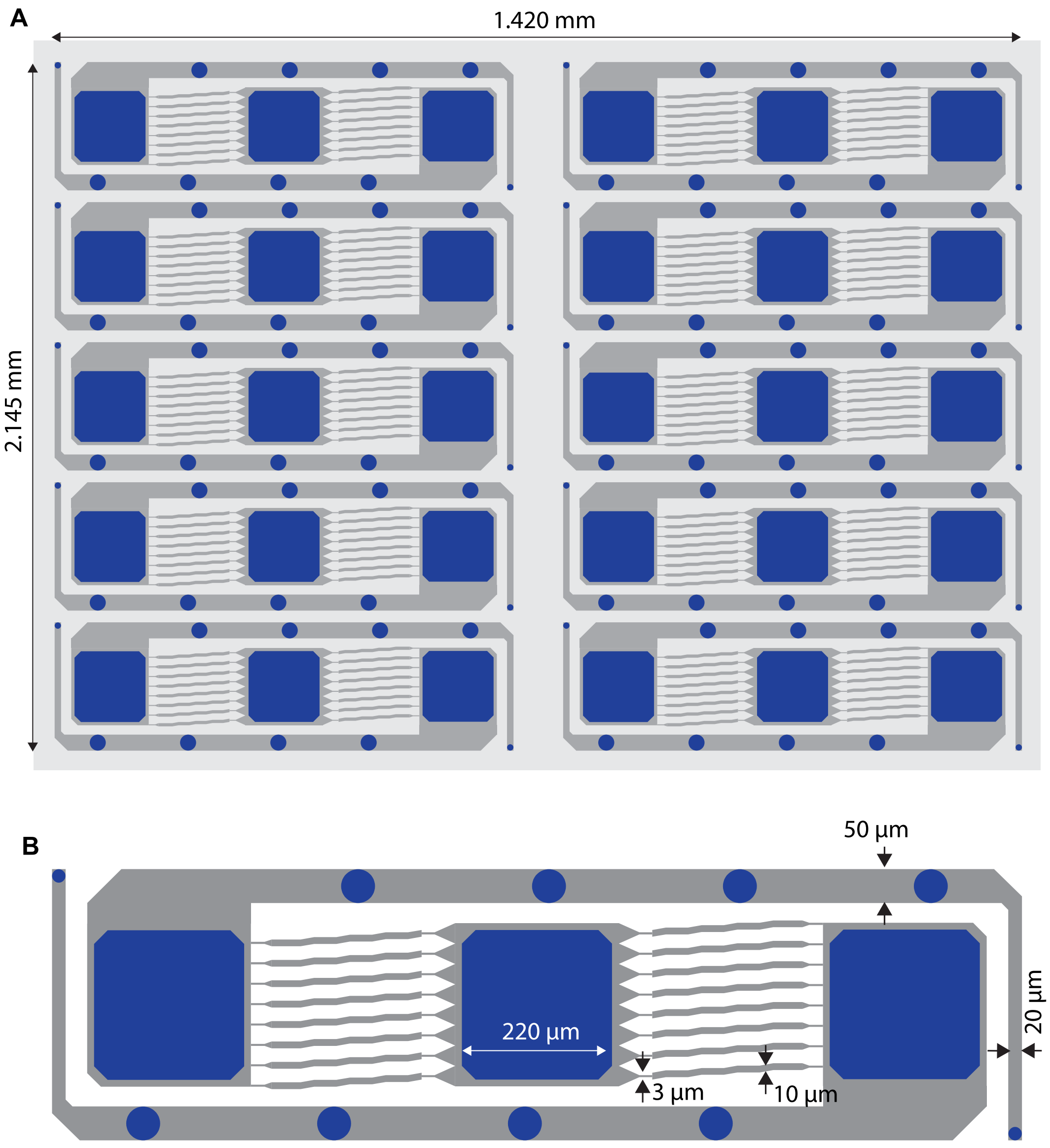


Figure S1: **Autocad design of the microstructure**. Each microstructure features 10 networks. Each network features a central seeding well for the hDRG spheroid. The lateral compartments allow better nutrient exchange and potential co-seeding of another cell type (e.g. fibroblasts). All networks are designed to optimize 1) Number of network per microstructure, 2) Number of microchannels per network, 3) Total number of covered electrodes per network (approx. 1000 electrodes which is about the maximum number of simultaneously recording electrodes on the chip), 4) usage of HD-MEA sensing area.

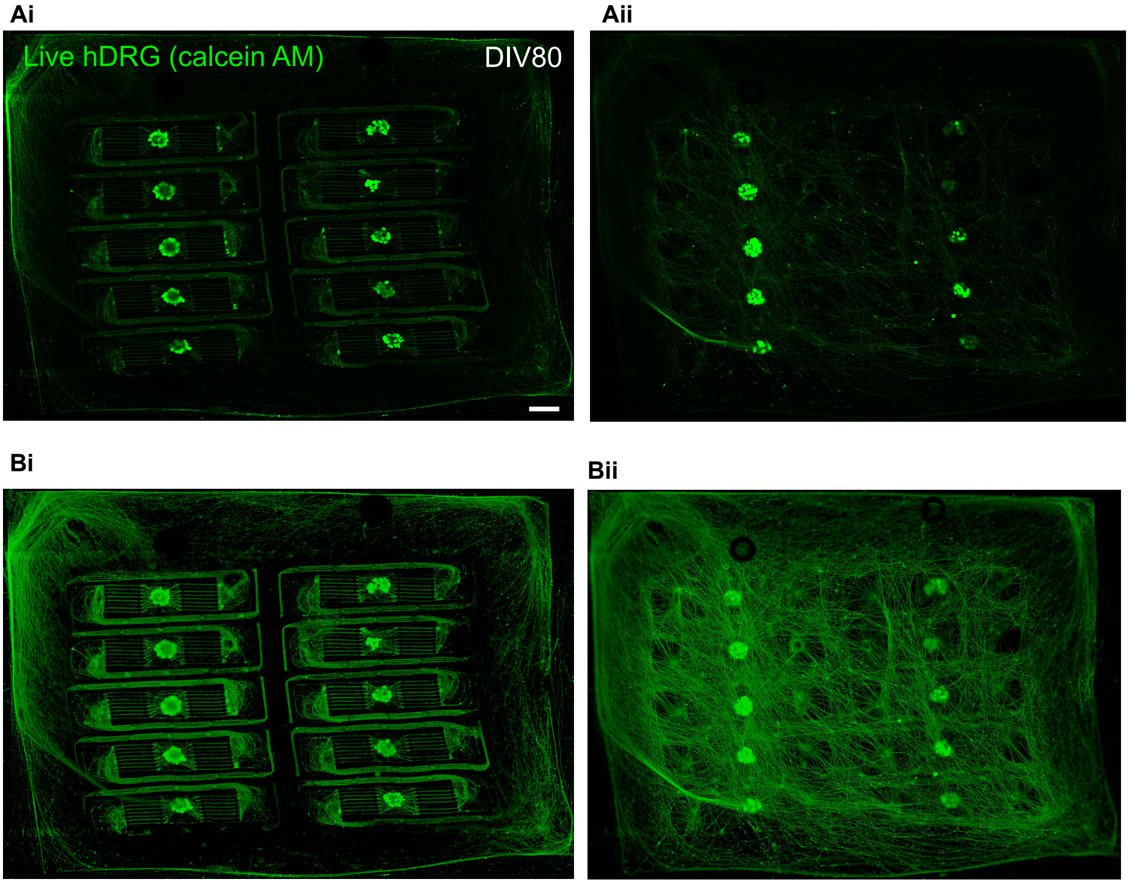


Figure S2: **Fluorescence staining of hDRG neuron cultures grown inside 10 networks at DIV80 on a HD-MEA chip. Calcein AM (green) stains live neurons. A**i Raw fluorescence image of the bottom plane (microchannel plane). Aii: image of the top of the microstructure (approximately 150 µm above). Axons grow on top due to the laminin coating everywhere, coming from the central seeding wells. **B** Log-processed image of A.

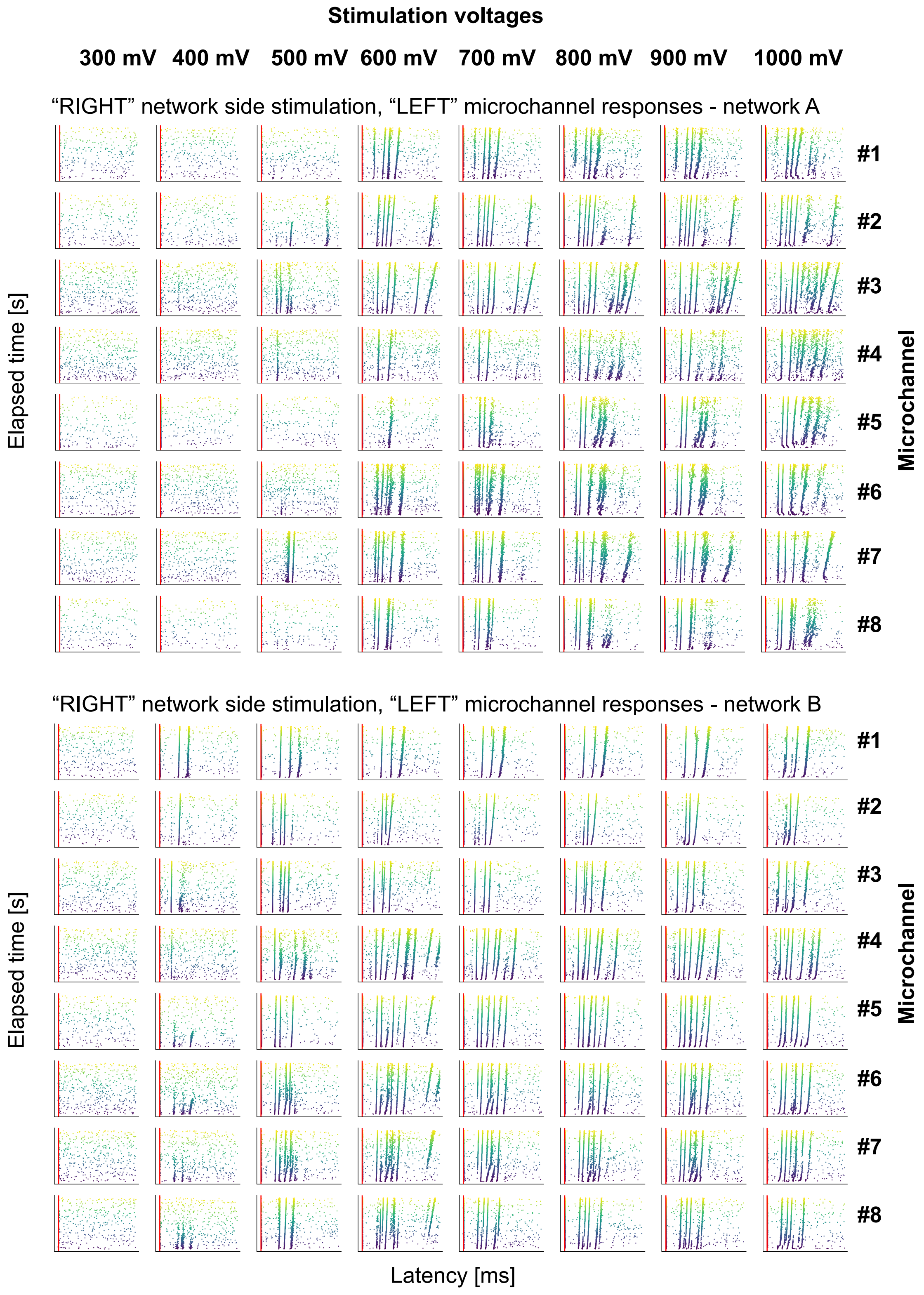


Figure S3: **Stimulation-latency response plot for all microchannels of two networks as example of voltage-dependent responses**

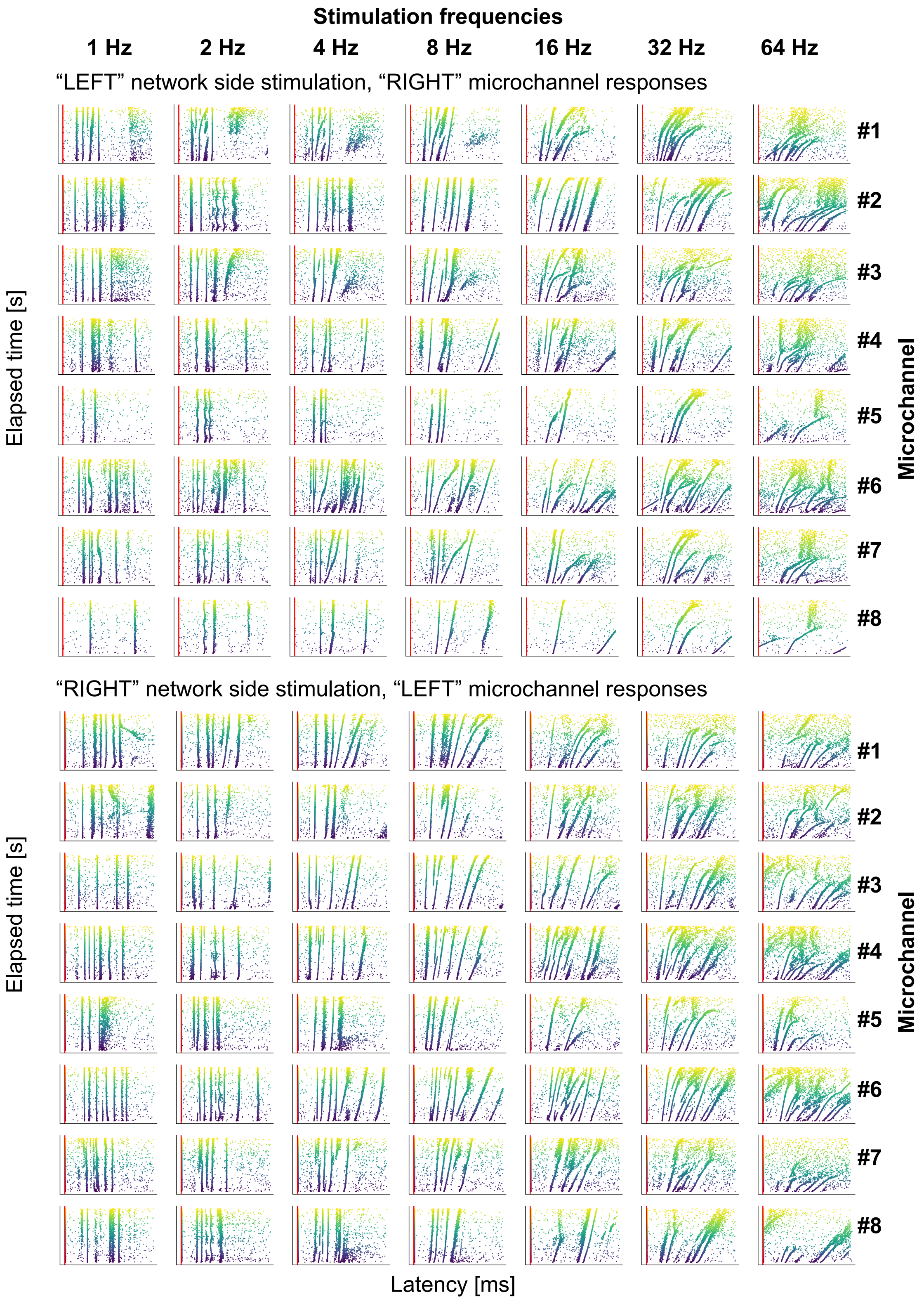


Figure S4: **Stimulation-latency response plot for all microchannels of two networks as example of frequency-dependent responses**

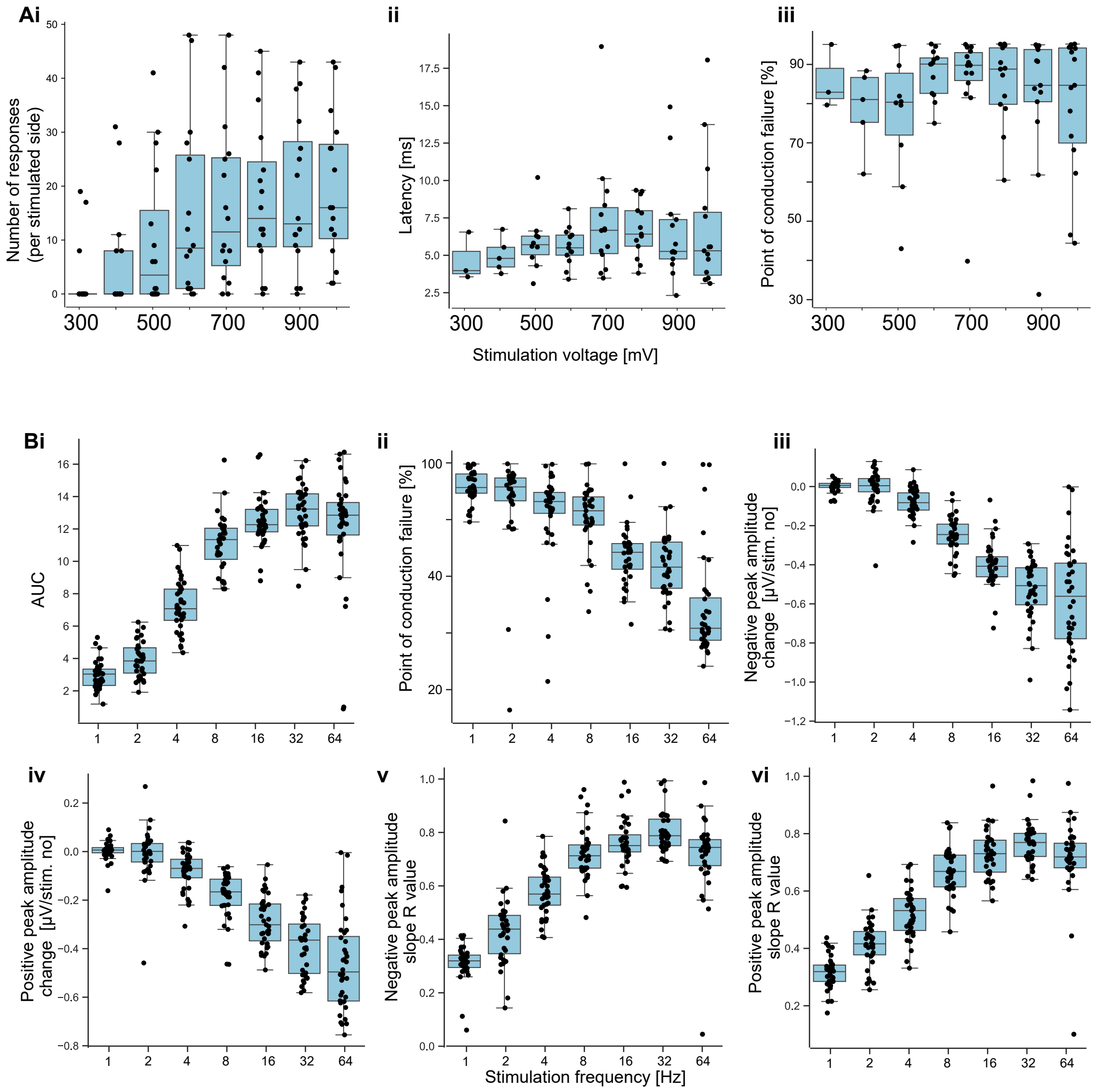


Figure S5: **A** Voltage-dependent network responses. i: Number of responses. ii: Latency. iii: Point of conduction failure. **B** Frequency-dependent responses. i: AUC. ii: Point of conduction failure. iii: Negative peak amplitude change. iv: Positive peak amplitude change. v: Negative peak amplitude slope R value. vi: Positive peak amplitude slope R value. All boxplots in B show the median (central line), the 25th and 75th (box edges), the whiskers represent data with 1.5x interquartile range. All datapoints are shown as individual points.

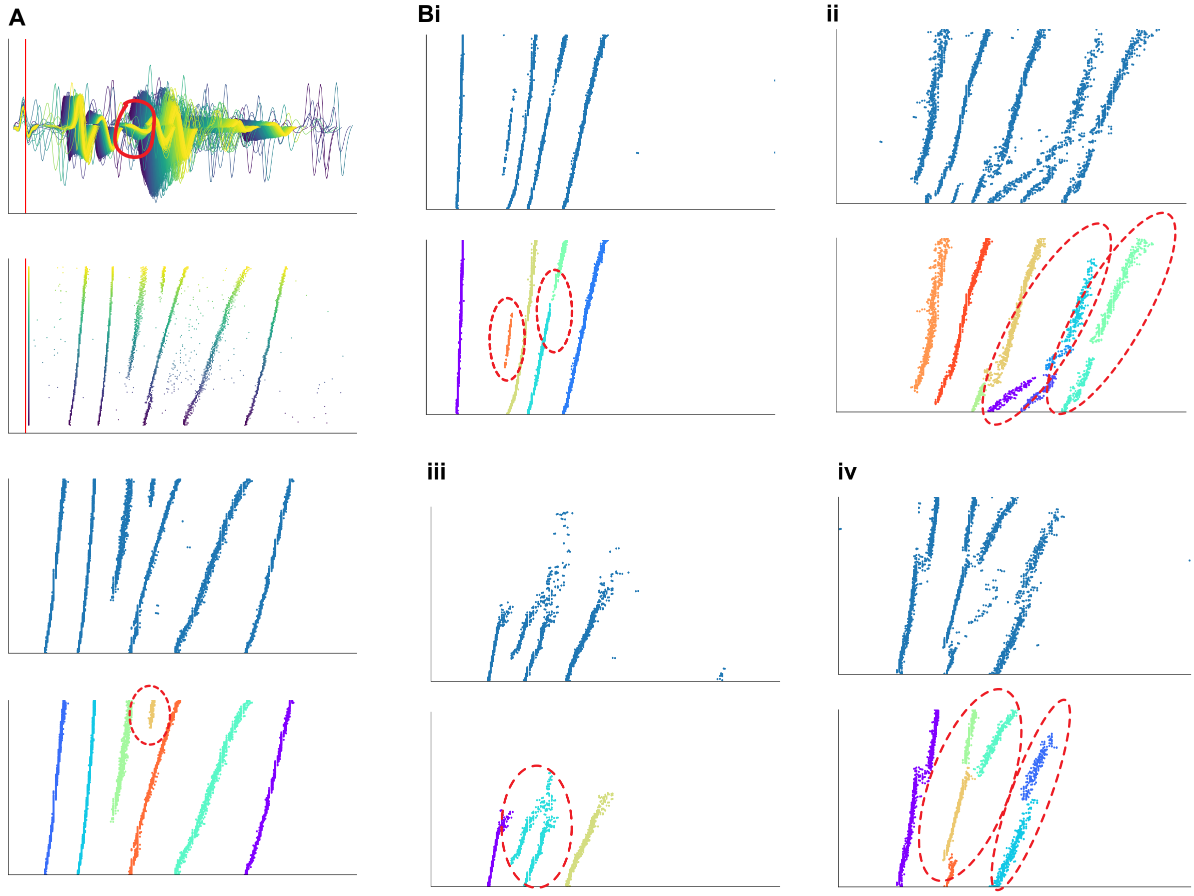


Figure S6: **Illustration of misidentified individual stimulation-induced response** Individual responses can be misidentified and sources are numerous, which impacts the accuracy of the metrics extraction. Case 1: due to the experimental set up, more than one axon grows per microchannel. We could have the case where two or more axons with very similar conduction speed and similar slow down rate would "compete" for detection (also result in more complex waveform) with overlapping latency which prevent the proper detection of one response (one axon) or the other. This induces some sudden shift back and forth in the band (see Biv). In this case, the peak with the largest amplitude was the detected one. Two side-by-side waveforms detection is limited by the 0.75 ms limit between two spikes. Case 2: in the case one of the two axons features a stronger slow down behaviour than the other one, a new "band" would appear later during the stimulation period (see A and Biv). This mainly impacts the initial latency metric. Case 3: due to the pre-processing pipeline and the detection algorithm, there is a filtering trade-off of filtering out spurious and spontaneous spikes forming background around bands and preserving the response integrity. If the filter is not strong enough, two close bands might be seen as a single one (see Biii). Due to the filtering, some small interruption in the response might be increased, leading to the opposite case: 2 identified bands instead of a single one (see Bi).

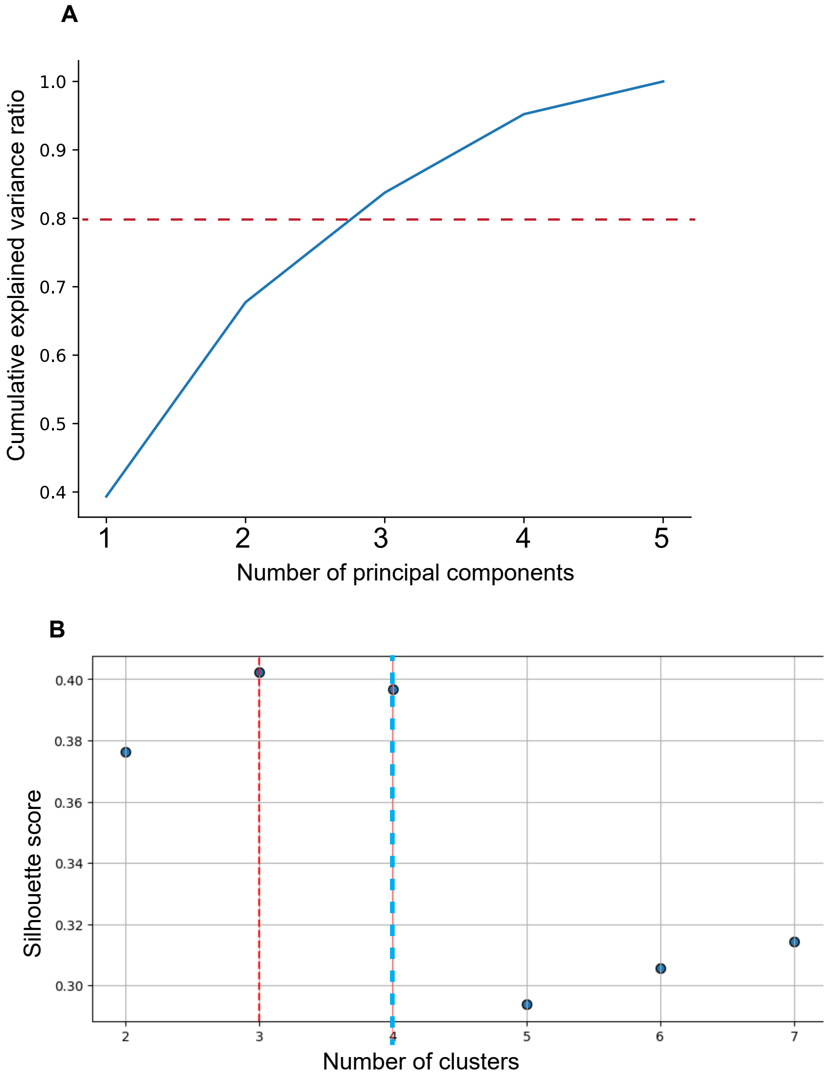


Figure S7: **PCA and cluster analysis. A**: Cumulative explained variance in function of number of principal components. Three principal components explain 80% of the total variance. **B**: The silhouette score was calculated for an increasing number of clusters inferred by Kmeans clustering.
